# Probing the functions of microglial cyclin-dependent kinase 5 under physiological and pathological conditions

**DOI:** 10.1101/2020.05.12.090605

**Authors:** Zhuyu Peng, Wen-Chin Huang, Maggie Chen, Jay Penney, Hugh Cam, Fatema Abdurrob, Leyla Akay, Xiao Chen, William T. Ralvenius, Lorena Pantano Rubino, Li-Huei Tsai

## Abstract

Cyclin dependent kinase 5 (Cdk5) regulates various developmental and physiological processes in the central nervous system. Deregulation of Cdk5 activity in neurons induces severe neurodegeneration and has been implicated in Alzheimer’s disease (AD) and other neurodegenerative conditions. A large fraction of AD risk genes are highly expressed in microglia, highlighting an important role for these cells in AD pathogenesis. While Cdk5 function in neurons is well characterized, our understanding of its roles in microglial function under physiological and neurodegenerative conditions remain rudimentary. Here, we investigate the roles of Cdk5 in microglia using myeloid-specific Cdk5 conditional knockout mice. Using microglia-specific transcriptome profiling, histological analyses, and behavioral assessments, we found that knockout of Cdk5 in microglia for 1 month induced transcriptional changes characterized by upregulation of cell cycle processes and type I interferon signaling genes in both physiological conditions and AD-related amyloidogenesis. In contrast to the robust transcriptional changes, conditional loss of microglial Cdk5 produced minimal effects on the density and morphology of microglia and their phagocytic activity toward myelin debris. Moreover, Cdk5cKO mice exhibited little change in synaptic density and tasks associated with locomotor, anxiety-like, and memory-related behaviors. Our findings indicate that the conditional loss of Cdk5 in microglia induces rapid alterations of microglial transcriptome with minimal or delayed effects on histological and behavioral responses.

## Introduction

Cyclin dependent kinase 5 (Cdk5) is a proline-directed serine/threonine kinase that primarily expressed in the central nervous system. Even though highly homologous in amino acid sequence to other CDK family members, Cdk5 is not involved in classical cell cycle regulation, in fact inhibiting the cell cycle in neurons^1,2^. Cdk5 also plays essential roles in regulating neuronal differentiation and migration, neurite outgrowth, axonal path finding, synaptic function and membrane transport^1–8^. Cdk5 null mice, which die near birth, exhibit severe defects in neuronal migration and inverted cerebral cortex laminar configurations^3^. Reconstitution of Cdk5 expression under control of the p35 promoter rescues the lethality phenotype in Cdk5 deficient mice^9^, indicating neuronal Cdk5 is vital for survival during embryogenesis and development^10^. Activation of Cdk5 requires association with one of its co-activators p35 or p39^11^,which exhibit temporal and spatial complementarity with Cdk5 in neurons in the CNS^12–14^. Although Cdk5/p35 complexes are essential in neuronal differentiation and migration, their roles in mature neurons, as well as in other brain cell types, remain less well understood.

In addition to its roles in neurodevelopment, Cdk5 activity has also been shown to be deregulated in various neurodegenerative diseases including Alzheimer’s Disease (AD)^15^, Parkinson’s Disease^16^ and amyotrophic lateral sclerosis^17^, with its pathological effects in AD being the best studied. AD is a pervasive neurodegenerative condition, characterized by progressive loss of memory and cognitive functions. Various pathological hallmarks have been well characterized, including β-amyloid deposition, hyperphosphorylation of the microtubule binding protein tau, loss of synapses and chronic neuroinflammation^18^. Physiological neuronal activity, as well as multiple neurodegenerative stimuli, lead to the calpain-mediated cleavage of Cdk5 activator p35 to a truncated form, p25^19^. p25 levels are found elevated in multiple neurodegenerative contexts and Cdk5-p25 complexes exhibit increased stability, distinct subcellular localization and altered substrate specificity compared to Cdk5-p35 complexes^20–24^. Consistently, Cdk5 activity is found elevated in postmortem AD patient brains as well as in AD models^25^ where deregulated Cdk5 contributes to hyperphosphorylation of amyloid precursor protein (APP)^26^, tau and neurofilament proteins^15,27^ and promotes neuronal death^15^. Importantly, inhibition of Cdk5 *in vivo* can attenuate tauopathy and amyloidogenesis and prevents neuronal loss and memory impairments^28–31^.

Growing evidence also indicates essential roles for Cdk5 in non-neuronal brain cell types^32–35^. Conditional deletion of Cdk5 in oligodendrocytes *in vivo* disrupts the integrity of nodes of Ranvier and significantly impairs mouse learning and memory^34^. *In vivo* and *in vitro* studies have further shown important roles for Cdk5 in promoting the transition of oligodendrocyte precursor cells to mature oligodendrocytes; conditional knockout of Cdk5 in Olig1^+^ cells results in failure in remyelination and reduced maturation of oligodendrocytes^35^.

The potential functions of Cdk5 in microglial cells have not yet been explored. Microglia are a specialized brain-resident macrophage population which arise from primitive hematopoietic precursors in the yolk sac and migrate to the CNS early in embryonic development^36^. Characterized by strong plasticity, microglia are capable of sensing various stimuli to produce appropriate cellular responses. Under physiological conditions microglia exhibit dynamic surveillance behavior by constantly extending and retracting their ramified processes to monitor the surrounding microenvironment. They are involved in refinement of brain circuits through complement-mediated synaptic pruning^37–39^, modulate myelin homeostasis^40–42^, scavenge apoptotic cellular debris^43,44^ and communicate with other cell types in the brain during resting states^43,45^. When CNS homeostasis is disrupted, resting microglia rapidly switch to activated phenotypes that can initiate immune responses, secrete inflammatory cytokines and chemokines, express cell surface antigens and exhibit altered morphologies in an attempt to sustain brain homeostasis^46,47^. However excessive increase of cytokines can lead to sustained activation of microglia, triggering a harmful positive feedback loop of inflammation^48^. Over the course of neurodegeneration in AD, microglia manifest heterogeneous transcriptional^49,50^, morphological and functional profiles in both humans^49^ and mice^50,51^. A single cell RNA-seq study demonstrated two distinct disease-associated microglial stages in the Ck-p25 mouse hippocampus^50^. At early stages, microglia show a proliferative transcriptional profile, while at later stages microglia acquire a signature characterized by distinct immune responses associated with type I and type II interferon signaling.

Another transcriptional study in 5xFAD mice showed a subset of microglia gradually adopting a disease-associated microglia (DAM) phenotype, whereby homeostatic microglial genes are first downregulated, followed by Trem2-dependent upregulation of DAM signatures, including genes relevant to phagocytosis and lipid metabolism^51^. Both studies support the view of a sequential transcriptional transformation of microglia subsets, indicating specific microglial signaling pathways are affected in a time- and context-dependent manner. Further investigation is necessary to determine the functional outcome of these transcriptional alterations.

Here, we investigate the roles of Cdk5 in microglia using myeloid-specific Cdk5 conditional knockout mice. By characterizing microglia specific transcriptomic profiles, histological, and behavioral phenotypes, we found that knockout of Cdk5 from microglia for one month induced considerable transcriptomic changes in both wild-type and 5XFAD backgrounds. Our analysis of microglial numbers, morphology, phagocytic activity and clustering, however, showed only minor effects of Cdk5 cKO on these parameters. Similarly, amyloid plaque accumulation in 5XFAD mice, and various measures of locomotor and behavioral performance in wild-type and 5XFAD mice, revealed little effect of Cdk5 deletion.

## Results

### Knockout of Cdk5 in microglia results in transcriptional changes

To begin to characterize the functions of Cdk5 in microglia, we crossed Cx3cr1-CreERT2 and Cdk5^f/f^ mice to generate *Cdk5* conditional knock out (Cdk5cKO) mice in which Cdk5 can be ablated in microglia upon tamoxifen (TAM) injection. This allowed us to investigate Cdk5 function in adult microglia, without affecting any potential functions during development. We injected 5-month-old mice with TAM or vehicle to generate Cdk5cKO and age matched control mice. After aging these mice for 30 days to 6 months of age, we collected and processed hippocampal samples for RNA sequencing (RNA-seq). Hippocampal homogenates from 4 control (Cx3cr1-CreERT2; Cdk5^f/f^ mice treated with vehicle) and 4 Cdk5cKO (Cx3cr1-CreERT2; Cdk5^f/f^ mice treated with TAM) mice were labeled with CD11b and CD45 then subjected to fluorescence activated cell sorting (FACS) to specifically isolate microglia (Figure 1A). Following RNA extraction and sequencing, we analyzed differentially expressed genes (DEGs) between conditions, identifying 465 downregulated genes and 269 upregulated genes in the microglia of Cdk5cKO mice compared to microglia from control mice (Figure 1B). As expected, *Cdk5* expression in Cdk5cKO microglia was reduced to undetectable levels (Figure 1C).

**Figure 1.**
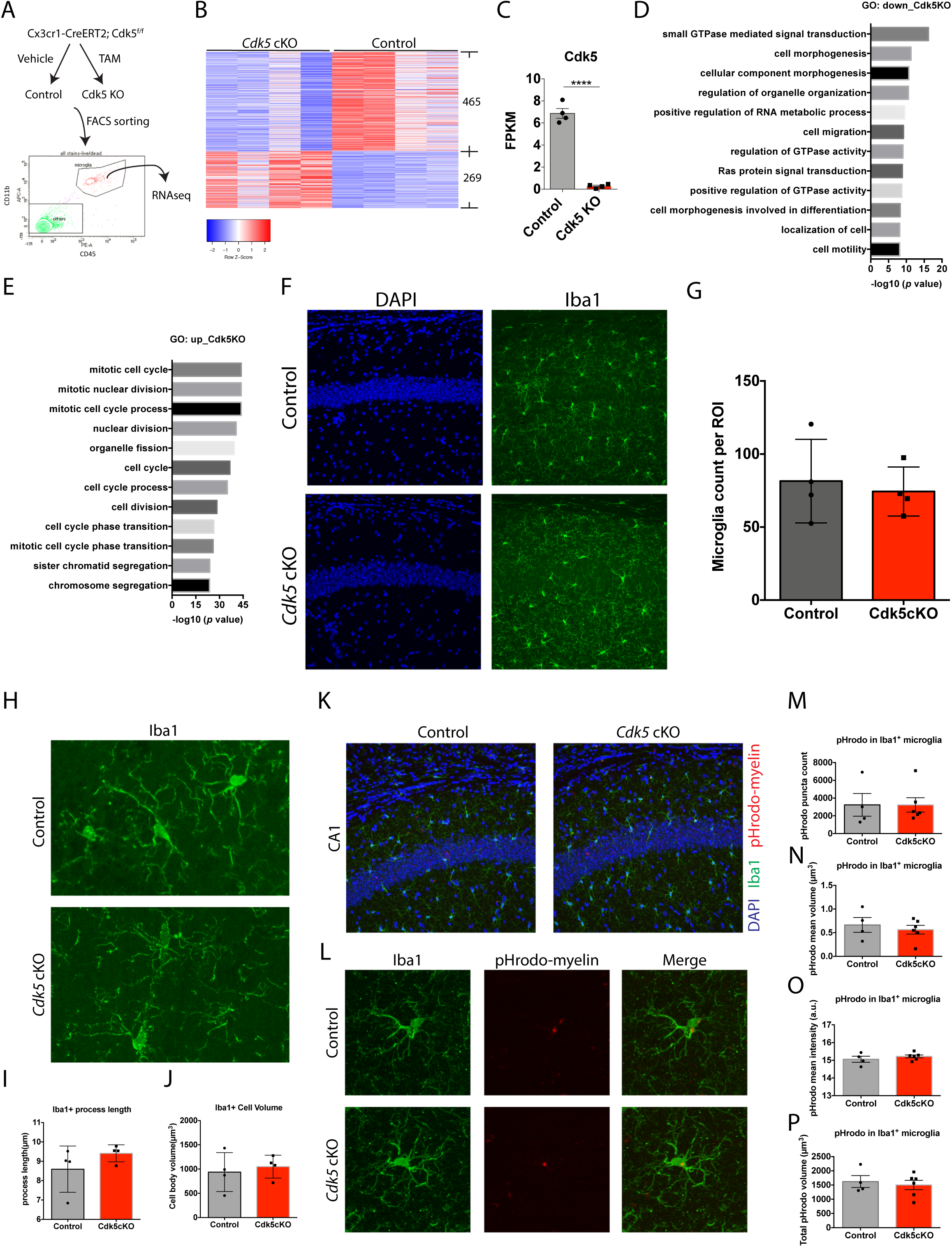
Knockout of *Cdk5* in microglia induces transcriptional changes. (A) A diagram showing the experimental flow of isolating microglia for bulk RNA sequencing. CD45 and CD11b were used to label microglia for fluorescence activated cell sorting. (B) A heatmap showing differentially expressed genes in microglia between control and Cdk5cKO mice. (C) Cdk5 expression was reduced in the RNA sequencing dataset. Unpaired t-tests were used, n=4 mice/samples per group. (D and E) Top GO biological process terms for genes downregulated (D) and upregulated (E) in Cdk5 KO microglia. (F) Images showing DAPI (blue) and Iba1 (green) in the CA1 region of the hippocampus. (G) Quantification of microglia numbers in the CA1 of the hippocampus in control and Cdk5cKO mice. Unpaired t-tests were used, n=4 mice per group. (H) Images showing Iba1^+^ (green) microglia morphology in the CA1 of the hippocampus. (I and J) Quantification of Iba1^+^ microglia process length (I) and cell body volume (J) in Cdk5cKO and control mice. Unpaired t-tests were used, n=4 mice per group. (K and L) Images showing pHrodo-myelin (red), Iba1 (green), and DAPI (blue) in the CA1 of the hippocampus (K) and magnified images (L). (M-P) Quantification of pHrodo-myelin signal in Iba1^+^ microglia. pHrodo puncta (M), mean volume (N), mean intensity (O), and total volume (P) were not significant different between groups. Unpaired t-tests were used, n=4 mice in control and n=6 mice in Cdk5cKO mice.

We next analyzed the DEGs we identified more carefully to better understand the potential functional outcomes of *Cdk5* KO in microglia. To this end, we performed Gene Ontology (GO) analysis to identify affected biological processes among the up- and downregulated genes. Genes downregulated in Cdk5cKO mice were enriched in GO terms associated with GTPase activity, cell migration, and cell motility (Figure 1D), while upregulated genes were enriched in GO terms associated with cell division and cell cycle processes (Figure 1E). These results suggest that microglial motility and proliferation may be altered in *Cdk5* KO microglia.

Having identified considerable gene expression changes in Cdk5cKO microglia, we then examined whether microglial density and morphology might be altered by loss of Cdk5. Thus, we performed histological analysis on another cohorts of mice, again one month after injection with either vehicle or TAM. Analysis of Iba1^+^ microglia in the CA1 region of the hippocampus showed that microglial density was not different between control and Cdk5cKO mice (Figure 1F and 1G). We also examined microglial morphology by measuring process length and cell body volume, again finding no significant differences between conditions (Figure 1H-J).

To assess whether phagocytic activity of microglia is altered in response to knockout of *Cdk5*, we next monitored *in vivo* phagocytic activity of microglia using a fluorescently labeled myelin uptake assay. We first purified myelin debris from C57BL/6 mice and conjugated it to pHrodo-Green dye, which is pH-sensitive and fluoresces at acidic pH. In biological systems, lysosomes are the main low-pH compartment, thus increased pHrodo-myelin fluorescence in this assay would reflect the final steps of cellular myelin uptake, trafficking through the endo-lysosomal system and deposition in acidic organelles. We injected pHrodo-myelin into the CA1 region of the hippocampus of 6-month-old control and Cdk5cKO mice (injected with vehicle or TAM one month prior) and perfused the mice 48 hours later for histological analysis. We observed pHrodo dye signal inside Iba1^+^ microglia in both conditions, indicating successful myelin uptake (Figure 1K and L). We next quantified the number, volume and average and total signal intensity of pHrodo-myelin puncta inside Iba1^+^ microglia, finding no evidence of altered myelin uptake by Cdk5cKO microglia (Figure 1M-P). Together, our findings indicate that knockout of *Cdk5* in microglia *in vivo* for one month induces considerable microglial transcriptional changes, but has minimal effect on the number, size, morphology, or phagocytic capacity of these cells.

### Knockout of Cdk5 in microglia does not affect synapse density or behavioral measures

Altered Cdk5 activity in neurons has profound effects on brain development, synaptic function and cognitive performance. To examine whether loss of Cdk5 in microglia also affects locomotor or cognitive function, we performed a battery of behavioral tests using control and Cdk5cKO mice. As for our earlier experiments, we used mice at 6 months of age that had been injected with vehicle or TAM one month prior. To assess gross locomotor activity, we performed the open field test and found that the total distance travelled by control and Cdk5cKO mice was similar (Figure 2A). We also measured the percent of time spent in the center of the open field arena, used as a measure of anxiety, and found no significant difference between control and Cdk5cKO mice (Figure 2B). To specifically probe anxiety-like behaviors, we further performed the light dark assay, in which a mouse was placed initially in the dark chamber then allowed to freely explore both dark and light chambers for ten minutes. A mouse spending more time in the light chamber is considered to be less anxious, as mice typically exhibit a preference for the dark. We found that both percent time spent in light chamber and latency to enter the light chamber were not significantly different between control and Cdk5cKO mice (Figure 2C and D), suggesting that anxiety-like behavior was not affected by loss of Cdk5. Next, to examine whether memory-related behaviors were affected in Cdk5cKO mice. we performed the novel object recognition test, comprising 3 phases over 4 days of testing. Each mouse was placed in an open field box for 10 minutes of habituation on day1, followed by two days of 10 minutes training on objection recognition. On the testing day (day 4), one of the two objects was replaced with a novel object, and each mouse was placed in the box to freely explore the two objects for 10 minutes. A discrimination index was calculated by dividing the time spent interacting with the novel object by the total time interacting with both the familiar and novel objects. We found that neither the discrimination index nor the time spent interacting with the novel object were significantly different between control and Cdk5cKO mice (Figure 2E and F).

**Figure 2.**
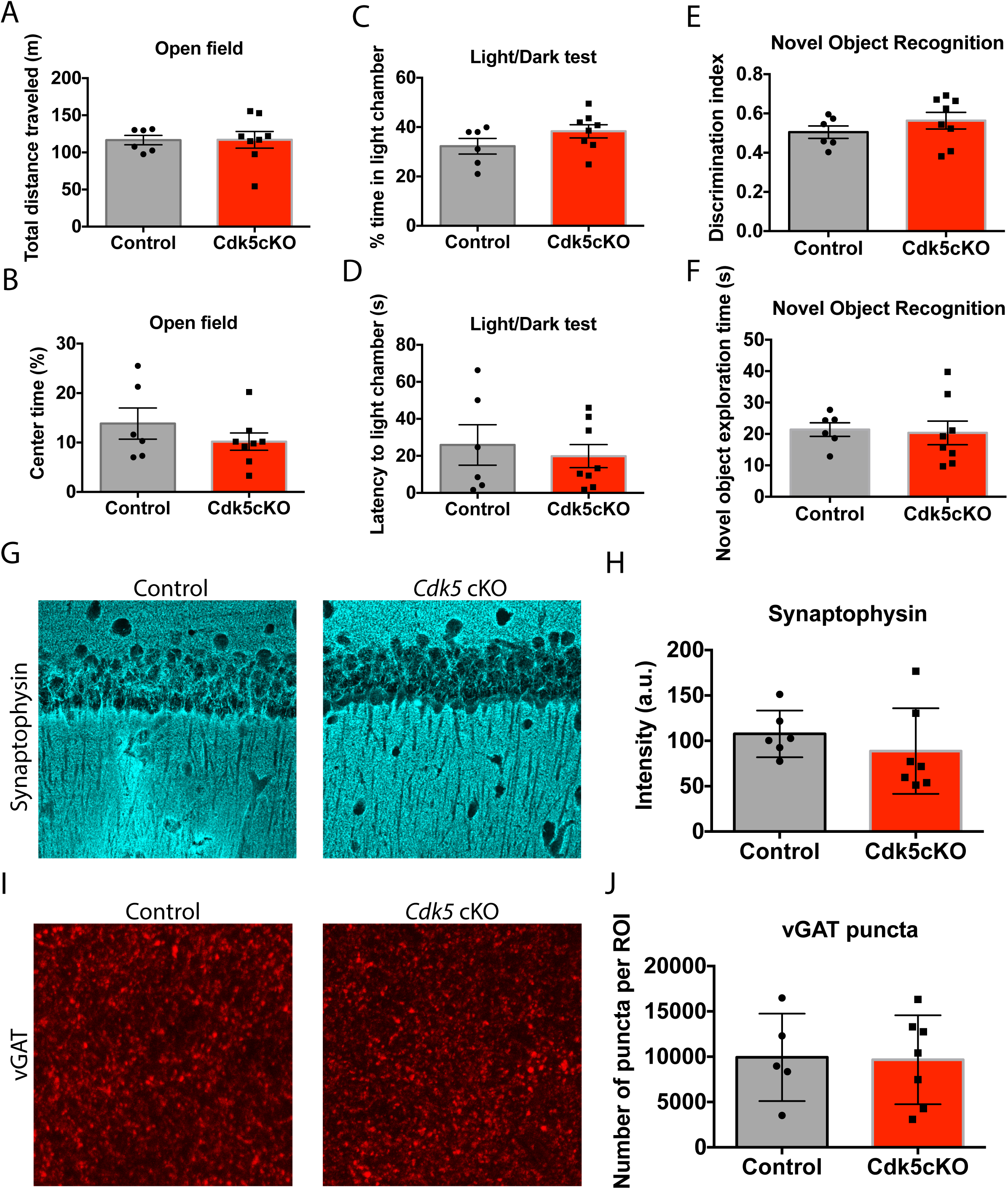
Knockout of *Cdk5* in microglia did not affect behavior or synaptic proteins. (A and B) Graphs showing total distance travelled (A) and percent time the mice spent in the center (B) in the open field test. Unpaired t-tests were used, n=6 mice in control and n=8 mice in Cdk5cKO mice. (C and D) Graphs showing percent time in light chamber (C) and latency to light chamber (D) in the light dark tests. Unpaired t-tests were used, n=6 mice in control and n=8 mice in Cdk5cKO mice. (E and F) Graphs showing discrimination index (E) and novel object exploration time (F) in the novel object recognition test. Unpaired t-tests were used, n=6 mice in control and n=8 mice in Cdk5cKO mice. (G and H) Images (G) and quantification (H) of synaptophysin signal intensity in the CA1 of the hippocampus. Unpaired t-tests were used, n=6 mice in control and n=7 mice in Cdk5cKO mice. (I and J) Images (I) and quantification (J) of vGAT puncta in the CA1 of the hippocampus. Unpaired t-tests were used, n=6 mice in control and n=7 mice in Cdk5 cKO mice.

We also examined markers of excitatory and inhibitory synapses in hippocampal area CA1 of control and Cdk5cKO mice. We examined the signal intensity of synaptophysin, a marker of glutamatergic synapses (Figure 2G and H) as well as quantifying the number of puncta associated with vesicular GABA transporter (vGAT), a marker of GABAergic synapses (Figure 2I and J). Consistent with the lack of effect we saw on behavioral measures in Cdk5cKO mice vs. controls, we did not observe any significant differences in these synaptic markers between conditions (Figure 2G-J).

### Loss of microglial Cdk5 in 5XFAD mice alters gene expression, proliferation and clustering

We next wanted to understand whether Cdk5 was important in microglia function in the context of AD, so we crossed Cx3cr1-CreERT2; Cdk5^f/f^ mice with 5XFAD mice to generate compound transgenic mice which we again injected with either vehicle (5XFAD; control) or TAM (5XFAD; Cdk5cKO). We first examined transcriptomic changes in response to knockout of Cdk5 in 5XFAD microglia as we had previously in control and Cdk5cKO mice. Vehicle or TAM were injected into 5-month-old mice, followed by FACS sorting to isolate microglia 30 days later, included 4 samples per group. Gene expression profiling and downstream analysis identified 409 upregulated genes and 203 downregulated genes in 5XFAD; Cdk5cKO microglia compared to those from 5XFAD; control mice (Figure 3A). Again, *Cdk5* itself was very strongly downregulated (Figure 3B). GO analysis revealed that downregulated genes were enriched in GO terms associated with cell activation, cytokine production, and cell motility (Figure 3C), while upregulated genes were enriched in GO terms associated with cell cycle processes, cell division, and innate immune responses (Figure 3D). Similar to our transcriptional findings in Cdk5cKO mice, compound mice also showed upregulation of genes associated with cell division and downregulation of genes associated with cell motility. To examine whether microglial density or morphology were altered in compound mice, we performed immunohistochemistry for Iba1. We found that knockout of Cdk5 in 5XFAD microglia resulted in increased Iba1^+^ microglia density in the CA1 region of the hippocampus (Figure 3E and F), while the morphology of these cells, as measured by cell body volume and process length, was not altered compared to controls (Figure 3G-I).

**Figure 3.**
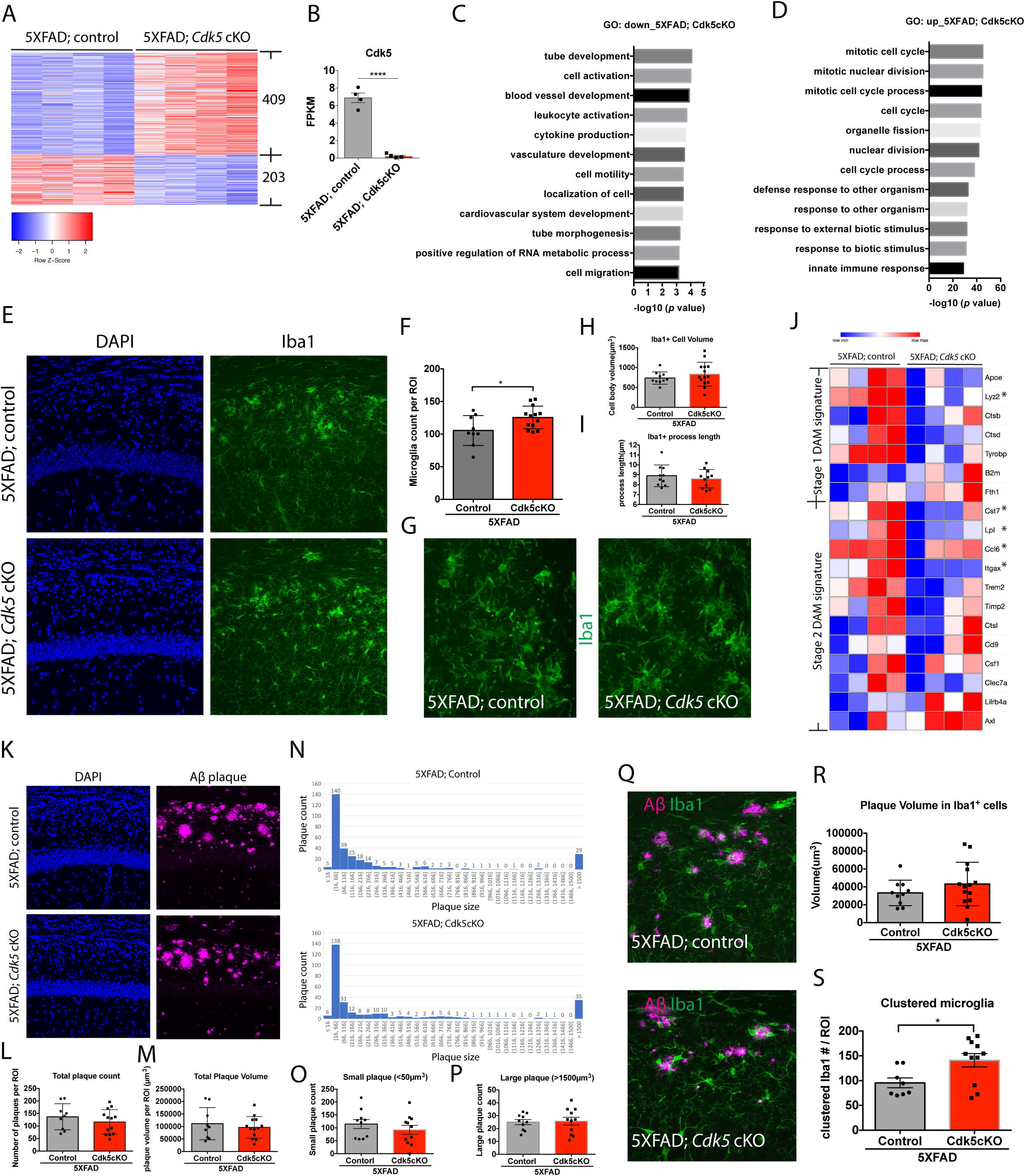
Profiling Cdk5 function in microglia of 5XFAD mice. (A) A heatmap showing differentially expressed genes between 5XFAD; control and 5XFAD; Cdk5cKO mice. (B) Cdk5 expression was significantly reduced in response to Cdk5 knockout in microglia in 5XFAD background. Unpaired t-tests were used, n=4 mice per group. (C and D) Top GO terms for genes downregulated (C) and upregulated (D) in 5XFAD; Cdk5cKO compared to 5XFAD; control mice. (E) Images showing DAPI (blue) and Iba1 (green) in the CA1 of the hippocampus in 5XFAD; Cdk5cKO and 5XFAD; control mice. (F) Quantification of Iba1^+^ microglia numbers in the CA1 of the hippocampus in 5XFAD; Cdk5cKO and 5XFAD; control mice. Unpaired t-tests were used, n=9 in 5XFAD; control and n=13 in 5XFAD; Cdk5cKO mice. (G) Images showing Iba1^+^ microglia morphology in 5XFAD; Cdk5cKO and 5XFAD; control mice. (H and I) Quantification of Iba1^+^ cell body volume (H) and process length (I) in 5XFAD; Cdk5cKO and 5XFAD; control mice. (J) A heatmap showing gene expression of stage I and II disease-associated microglia (DAM) signatures in 5XFAD; Cdk5cKO and 5XFAD; control mice. * indicate statistically significant genes. (K) Images showing DAPI (blue) and Aβ (magenta) in the CA1 of the hippocampus. (L and M) Quantification of plaque count (L) and volume (M) in the CA1 of the hippocampus. Unpaired t-tests were used, n=9 in 5XFAD; control and n=13 in 5XFAD; Cdk5cKO mice. (N) Graphs showing examples of plaque distribution in 5XFAD; Cdk5cKO and 5XFAD; control mice. (O and P) Quantification of small plaques (O) and large plaques (P) in the CA1 of the hippocampus. Small plaque was defined by <50µm^3^ in size and large plaque was defined by >1500µm^3^ in size. Unpaired t-tests were used, n=10 in 5XFAD; control and n=11 in 5XFAD; Cdk5cKO mice. (Q) Images showing Aβ (magenta) and Iba1 (green) in the CA1 of the hippocampus in 5XFAD; Cdk5cKO and 5XFAD; control mice. (R and S) Quantification of plaque volume in Iba1^+^ cells (R) and of clustered microglia (S) around plaques. Unpaired t-tests were used, n=10 in 5XFAD; control and n=14 in 5XFAD; Cdk5cKO mice (R). n=8 in 5XFAD; control and n=11 in 5XFAD; Cdk5cKO mice (S).

Previous studies have identified subsets of microglia associated with amyloid pathology^51^ in 5XFAD mice. These microglia were termed disease-associated microglia (DAM) and characterized by upregulation of a specific subset of genes. We examined the expression of these genes to understand whether knockout of Cdk5 from microglia in 5XFAD background would impact the expression of the DAM signature. Indeed, we found that many of the genes comprising both the stage 1 and stage 2 DAM signatures were expressed at lower levels in microglia from compound mice compared to those from 5XFAD; control mice, though not all reached statistical significance (Figure 3J).

Given that DAM microglia are associated with amyloid plaques and many DAM signature genes were downregulated in compound mice, we further examined amyloid burden in these mice. We performed immunohistochemistry for Aβ and quantified the number and volumes of amyloid plaques in the CA1 region of the hippocampus. We quantified plaques that were at least 5µm^2^ or bigger and found that both the plaque count and total plaque volume were comparable between 5XFAD; control and compound mice (Figure 3K-M). To understand if small and large plaques could be differentially affected by the loss of Cdk5 in 5XFAD mice, we quantified the number of small (<50µm^2^) and large plaques (>1500µm^2^). We found that regardless of size, the number of plaques was similar between control and compound mice (Figure 3 N-P). Thus, the number and composition of amyloid plaques in 6-month old 5XFAD mice does not appear to be altered by loss of microglial Cdk5.

Although amyloid plaques in 5XFAD mice were not altered by the conditional loss of Cdk5, the number of microglia was increased and the expression of DAM signature genes was reduced. This result prompted us to investigate whether phagocytic activity of microglia toward Aβ was changed in compound mice. We performed immunohistochemistry simultaneously for Iba1 and Aβ and quantified plaque volumes inside Iba1^+^ microglia (Figure 3Q). While the plaque volumes inside Iba1^+^ regions were not different between conditions (Figure 3R), the total number of microglia clustered around amyloid plaques, a classical feature of DAM microglia, was increased in compound mice compared to control (Figure 3S).

To understand whether knockout of Cdk5 in microglia could affect behavioral phenotypes in 5XFAD mice, we then performed open field, light/dark, and novel object recognition tests to probe locomotor, anxiety-like, and memory-related performance. We found that in the open field test, both total distance traveled and percent time in center were not altered between groups (Figure 4A and B). In the light/dark test, although percent time spent in the light chamber remained similar between the two groups (Figure 4C), compound mice showed slightly increased latency to the light chamber (Figure 4D). In the novel object recognition test, both discrimination index and time spent exploring the novel object were not significantly different between the two groups (Figure 4E and F). Together, these results suggest that while knockout of Cdk5 from microglia in a 5XFAD background affects microglial transcription, abundance and clustering, it has minimal effects on amyloid deposition or behavioral performance.

**Figure 4.**
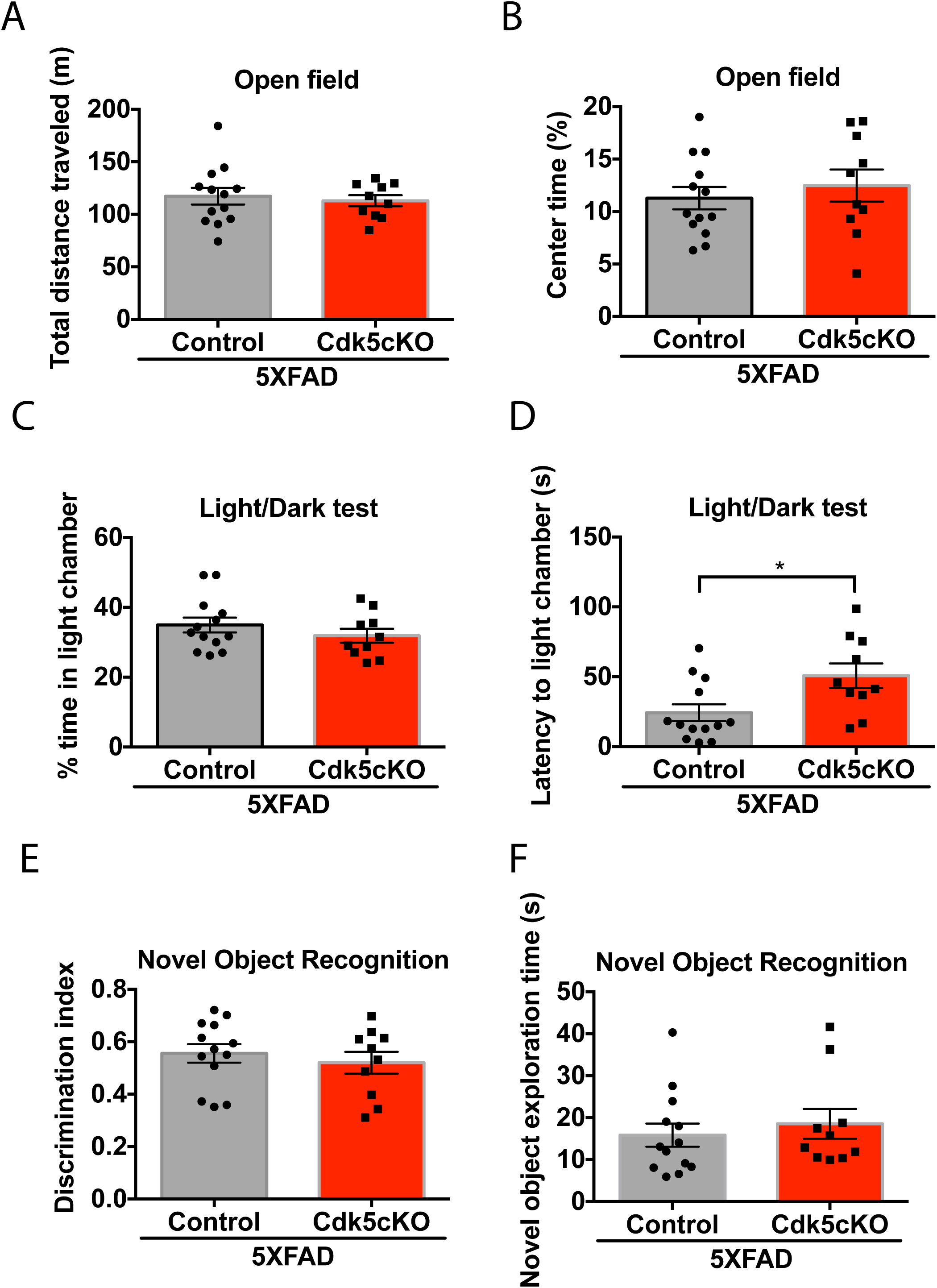
Behavioral characterization of 5XFAD mice in response to Cdk5 knockout in microglia. (A and B) Graphs showing total distance travelled (A) and percent time the mice spent in the center (B) in the open field test. (C and D) Graphs showing percent time in light chamber (C) and latency to light chamber (D) in the light dark tests. (E and F) Graphs showing discrimination index (E) and novel object exploration time (F) in the novel object recognition tests. Unpaired t-tests were used, n=13 mice in 5XFAD; control and n=10 mice in 5XFAD; Cdk5cKO mice.

### Knockout of Cdk5 from microglia upregulates cell cycle and type I interferon signaling genes across physiological and pathological states

We next wanted to examine genes and biological processes that are dysregulated in response to loss of *Cdk5* from microglia in both wild-type and 5XFAD backgrounds. We found that of the 465 downregulated genes in Cdk5cKO vs. control microglia, 71 genes were also downregulated in compound vs. 5XFAD; control microglia (Figure 5A). Of the 269 upregulated genes in Cdk5cKO vs. control microglia, 158 genes were also upregulated in compound vs. 5XFAD; control microglia (Figure 5A). Thus, a considerable portion of the Cdk5-dysregulated genes were affected in microglia regardless of genetic backgrounds (wild-type vs. 5XFAD). GO analysis of these commonly dysregulated genes revealed that Cdk5-downregulated genes were enriched in GO terms associated with regulation of lipid transport and transcription (Figure 5B), and that commonly upregulated genes were enriched in GO terms associated with cell cycle processes and type I interferon signaling (Figure 5C).

**Figure 5.**
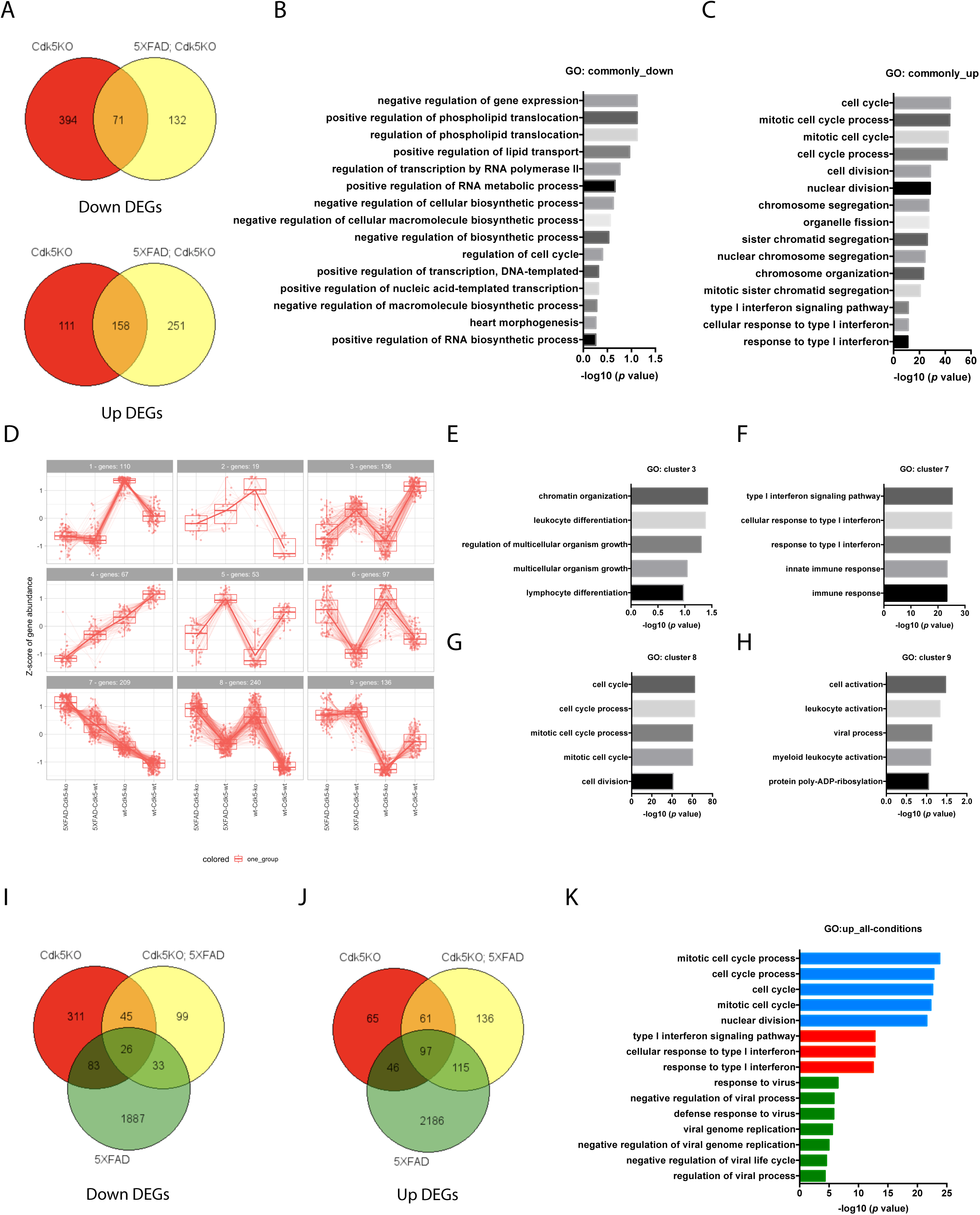
Transcriptomic analysis of commonly dysregulated genes in response to knockout of Cdk5 in microglia. (A)Venn diagram showing commonly downregulated and upregulated genes between control vs Cdk5cKO and 5XFAD; control vs 5XFAD; Cdk5cKO microglia. (B and C) Top enriched GO terms for commonly downregulated (B) and commonly upregulated (C) genes in control vs Cdk5cKO and 5XFAD; control vs 5XFAD; Cdk5cKO comparisons. (D) Gene expression patterns of differentially expressed genes between control vs Cdk5cKO and 5XFAD; control vs 5XFAD; Cdk5cKO. Data were plotted in 4 genotypes, including wt-Cdk5-wt, wt-Cdk5-ko, 5XFAD-Cdk5-wt, and 5XFAD-Cdk5-ko. 9 clusters of gene expression patterns were identified. (E-H) Top enriched GO terms for genes in cluster 3 (E), 7 (F), 8 (G), and 9 (H). (I and J) Venn diagrams showing commonly downregulated (I) and commonly upregulated (J) genes in the comparisons among control vs Cdk5cKO, 5XFAD; control vs 5XFAD; Cdk5cKO, and control vs 5XFAD; control. (K) Selective enriched GO terms for commonly upregulated genes across the three comparisons.

To understand how these genes and biological processes were dysregulated across the four genotypes (control, Cdk5cKO, 5XFAD and compound), we analyzed and plotted the expression patterns of DEGs across the four genotypes, identifying nine clusters of expression patterns (Figure 5D). While some clusters contained very few genes, we focused on the four clusters that contained >100 genes, performing GO analysis on the gene lists from these clusters. The genes in cluster 3, which showed slight downregulation in 5XFAD compared to control microglia and showed further downregulation in cells from Cdk5cKO and compound mice, were enriched in GO terms associated with chromatin organization and leukocyte differentiation (Figure 5E). The genes in cluster 7, which showed gradual upregulation across all four genotypes, were enriched in GO terms associated with type I interferon signaling (Figure 5F). The genes in cluster 8, which exhibited slight upregulation in 5XFAD compared to control mice and further upregulation in Cdk5cKO and compound mice, were enriched in GO terms associated with cell cycle processes (Figure 5G). The genes in cluster 9, which showed upregulation in 5XFAD compared to control mice and showed downregulation in Cdk5cKO mice but no change in compound mice compared to 5XFAD mice were enriched for cell activation (Figure 5H).

We further performed GO analysis to uncover genes and pathways that were commonly dysregulated across different pairwise groups (control vs 5XFAD, control vs Cdk5cKO, and 5XFAD vs compound). We found 26 downregulated genes and 97 upregulated genes shared across all three pairwise groups (Figure 5I and J). While there were no enriched GO terms for the commonly downregulated genes, the commonly upregulated genes were enriched in GO terms associated with cell cycle processes, type I interferon signaling, and defense response to viruses (Figure 5K). Together, our results indicate that the most robust gene dysregulation signatures in response to conditional loss of Cdk5 in microglia were upregulation of cell division, type I interferon signaling, and defense response to viruses.

## Discussion

A large fraction of AD risk genes are highly expressed in microglia, implicating these cells in the pathogenesis of AD^**52–54**^. Dysregulated Cdk5 activity has also been observed in AD and other neurodegenerative conditions^**15–17**^, while animal models have demonstrated the damaging effects of Cdk5 loss^**3**^ or hyperactivity^**19**^. Thus, understanding the roles of Cdk5 in microglia and the potential effects on AD pathogenesis are of considerable interest. Using transcriptomic, histopathological, and behavioral assays, we profiled the role of Cdk5 in microglia *in vivo*. We found that conditional knockout of Cdk5 from microglia of either wild-type or 5XFAD mice induced robust transcriptional changes, including upregulation of cell cycle processes, type I interferon signaling, and defense response to viruses. Despite the gene expression changes we observed, we found Cdk5cKO had only modest effects at the levels of cellular composition and morphology as well as on performance tasks that assess locomotor, anxiety, and memory functions. The lack of strong phenotypic changes beyond altered gene expression could reflect homeostatic mechanisms that prevent or obstruct responses at the cellular and behavioral levels. It is also possible that deletion of microglial Cdk5 for longer than one month, as was performed here, is required for the manifestation of phenotypic changes.

Knockout of *Cdk5* in microglia for one month induced expression changes for hundreds of genes in both the wild-type and 5XFAD backgrounds. Interestingly, many of the upregulated genes were also upregulated in the microglia of 5XFAD compared to control mice, specifically genes associated with cell division, type I interferon signaling, and defense response to viruses. This suggests that knockout of *Cdk5* in microglia recapitulates some aspects of microglial disfunction in 5XFAD mice. In contrast, DAM signature genes, which are upregulated in 5XFAD compared to control mice, were downregulated in response to knockout of *Cdk5* in the microglia of 5XFAD mice, suggesting Cdk5cKO may allow microglia in 5XFAD brains to maintain a more homeostatic state. DAM microglia are also known to cluster around amyloid plaques^51^, but whether they contribute to reducing amyloid load is not clear. Our findings indicate that while DAM signature genes were downregulated in microglia upon loss of *Cdk5*, the amyloid burden was not altered, suggesting that there is no direct correlation between upregulation of DAM signature genes and amyloid burden. It is also possible that cKO of microglial Cdk5 for a longer period may be necessary to observe effects on amyloid load in 5XFAD mice.

Although we observed transcriptomic changes in microglia in response to knockout of *Cdk5*, we did not observe clear histological changes as suggested by our transcriptomic findings. This indicates that gene expression changes in response to knockout of *Cdk5* in microglia only have minimal effects on histological phenotypes we examined in this study. Alternatively, given that we examined both transcriptomic changes and the histological phenotypes one month after knockout of *Cdk5* (via TMA injections), alteration of histological phenotypes may lag behind transcriptional changes. Future experiments examining the histological phenotypes at later time point may resolve these two possibilities.

## Acknowledgements

We thank all the members of the Tsai laboratory for technical assistance and comments on this manuscript. This research was supported in part by the JBP foundation and National Institute of Health (NIH) grants R37NS051874 to L-HT.

## Author Contributions

WCH, ZP, and L-HT conceived the study. WCH and ZP designed the study. ZP, MG, FA, XC, LA, WR performed experiments and analyzed data. WCH, HC, and LPR analyzed the RNA sequencing data. WCH, ZP, JP, and HC wrote the paper with inputs from all the authors.

## Method

### Mice

Cx3Cr1-CreERT2 and 5XFAD mice were obtained from the Jackson Laboratory and Cdk5^f/f^ mice were produced previous by the Tsai lab^55^. Tamoxifen or vehicle were injected in 5-month-old mice and transcriptomic, histological, and behavioral assays were carried out at 6 months of age. Male mice were used in this study. All procedures were conducted in accordance with the U.S. NIH Guide for the Care and Use of Laboratory Animals and were approved by The Committee for Animal Care of the Division of Comparative Medicine at the Massachusetts Institute of Technology.

### Immunohistochemistry

Mouse brains were fixed in 4% PFA for one night and sectioned coronally at 40 µm using a vibrotome (Leica). Brain sections were washed with PBS solution for 10 minutes and incubated in blocking solution (1% BSA in 0.3% Triton X-100, PBS) for two hours at room temperature. The sections were incubated in primary antibodies in blocking solution for 48h at 4°C. Primary antibodies used in this study include anti-β-amyloid (D54D2)(1:500; Cell Signaling Technology; 8243), anti-Iba1 (1:500; Synaptic Systems 234 004, polyclonal guinea pig anti-serum), anti-Synaptophysin antibody (1:500; Abcam ab32137) and VGAT (1:500; Synaptic Systems 131 013). The sections were washed with PBS solution (4 × 10 min) and incubated in secondary antibodies (dilution, 1:2,000) in blocking solution for one night at 4°C. Primary antibodies were visualized by using Alexa – Fluor -488, Alexa – Fluor -594, and Alexa – Fluor -647 antibodies (Life Technologies). All cell nuclei were visualized with Hoechst 33342 (Thermo Fisher Scientific, #H3570). Sections were washed with PBS (4 × 10 min) before being mounted with Prolong Gold mounting medium (Thermo Fisher Scientific, #P36930). Images were obtained with Zeiss confocal microscope LSM 710 or LSM 880. For each experimental condition, two images were acquired for quantification.

### Microglia-specific RNA sequencing

Hippocampal tissues were rapidly removed from mouse brains, placed in ice-cold Hanks’ balanced salt solution (HBSS) (Gibco by Life Technologies, catalogue number 14175-095) and cell suspensions were resolved by enzymatic digestion using the Neural Tissue Dissociation Kit (P) (Miltenyi Biotec, catalogue number 130-092-628). Cell suspension was then stained using allophycocyanin (APC)-conjugated CD11b mouse clone M1/70.15.11.5 (Miltenyi Biotec, 130-098-088) and phycoerythrin (PE)-conjugated CD45 antibody (BD Pharmingen, 553081). FACS was then used to purify CD11b and CD45 positive microglial cells. BD FACSDiva 8.0 and FlowJo V10 (Tree Star, Inc.) were used to for FACS analysis. Standard side scatter width versus area and forward scatter width versus area criteria were used to discriminate doublets and gate only singlets. Viable cells were identified by staining with propidium iodide (PI) and gating only PI-negative cells. RNA was extracted using RNeasy Plus Mini Kit (QIAGEN, 74134) according to the manufacturer’s protocol, and total RNA was treated with DNase I (Worthington Biochemical) followed by RNA purification using RNA Clean and Concentrator-5 Kit (Zymo Research) according to manufacturer’s protocols. Purified RNA was subjected to quality control (Fragment Analyzer). cDNA libraries were prepared using Illumina TruSeq Total RNA Sample Prep Kits (Illumina) and High-Throughput 3’Digital Gene Expression (HT-3’DGE). Libraries were sequenced on the Illumina HiSeq 2000 platform at MIT BioMicro Center. The collapsed raw fastq reads were aligned by Tophat2, and further processed by Cufflinks 2.0.0 with UCSC mm9 reference gene annotation to determine transcript abundance. A gene was considered differentially expressed with a fold change ≥ 1.5 and a statistical significance of P< 0.05. Gene ontology (GO) analysis of DEGs were performed using ToppGene.

### Microglia analysis

#### Analysis of microglia number

3D rendering and quantification was performed by Imarisx64 8.1.2 (Bitplane, Zurich, Switzerland). Imaris spot module allowed placement of a sphere at the center of the soma of each cell based on the Iba1 and Hoechst signals, thereby counting the Iba1-positive microglia number.

#### Analysis of microglia morphology

Individual microglia branch lengths were measured using Imaris filament function. Filament tracer function traced the start points and end points of microglia processes and presented associated statistics.

#### Microglia clustering analysis

Microglia clustering pattern around amyloid plaques was analyzed in 40um slices. Imaris detected D54D2 signals and rendered amyloid plaque surfaces accordingly. Spot module was implemented for counting Iba1-positive microglia number by placing a spot at the center of the soma of each cell. And then the Spots Close to Surface XTension function counted spots within 25µm radius to the amyloid plaques surfaces and excluded the spots outside of this range.

### Microglia phagocytic activity toward myelin debris

#### Myelin extraction

Brains were homogenized with a douncer containing 0.32 M sucrose and subsequently layered on 0.85 M sucrose and ultracentrifuged using a SW32Ti rotor spun at 75 000xg for 40 min. The interphase was then exposed to osmotic shock by twice-repeated resuspension in distilled water followed by ultracentrifugation at 75 000xg for 15 min. Then the pellet was resuspended in 0.32 M sucrose and layered on 0.85 M sucrose, spun at 75 000xg for 30 min and the interphase was washed twice in distilled water and finally resuspended in Tris-buffered saline.

#### pHrodo-Myelin preparation

1 mg pHrodo Green STP Ester (amine-reactive) was dissolved in 150 µL DMSO and then added to Eppendorf tubes with approximately 20 mg myelin. Then the solution was left in darkness for 30 min before adding to 10 mL DPBS. Solution was spun at 12 000 xg (9 800 rpm) for 15 min. Next, supernatant was discarded and the pellet was resuspended in 10mL DPBS, then spun at 12 000 xg (9 800 rpm) for another 10 min. The supernatant was then discarded and washed. Finally, the pellet was resuspended in DPBS at 1 mg/mL protein (from Nanodrop, 280A) and preserved at −20°C.

#### Stereotaxic injection of pHrodo-Myelin

To characterize the microglia phagotcytic behavior in vivo, stereotaxic injection of 500nL pHrodo-Myelin (1mg/ml) was injected to the CA1 (A|P: −2.25mm, M|L: 1.75mm, D|V: −1.5mm relative to the Bregma), with an infusion rate at 100nL per minute. The needle stayed at the target injection coordinate for 3 minutes before the infusion and for 5 minutes after injection was complete.

#### Analyses of pHrodo-Myelin

Imaris Surfaces function was utilized detect and 3D render Iba1 signals in 3D space. Then the mask function was then implemented to characterize the colocalization of pHrodo-myelin distribution within microglia surfaces. The mask feature allowed us to ignore signal intensity outside of region of interest and generate a new channel containing only voxels in the region of interest, using the pHrodo-myelin channel as the source channel. Imaris Surfaces function were utilized again to detect and 3D render D54D2 signals in this new channel, allowing for characterization of pHrodo-myelin colocalization with microglial cells in 3D space.

### Synapse density analysis

Imaris Surfaces function was utilized to detect and 3D render vGAT and Iba1 signals respectively, with associated statistics presented. Mean signal intensity of synaptophysin was measured using Image J.

### Analysis of amyloid plaques

Amyloid beta plaques were detected and rendered with Imaris surfaces module based on D54D2 signals, with associated statistics including the number of rendered surfaces and volume of each individual surface presented.

#### Plaque colocalization with microglia

Imaris Surfaces function was utilized to detect and 3D render Iba1 signals in 3D space. Then the mask function was implemented to characterize the colocalization of amyloid plaque distribution within microglia surfaces. The mask feature allows to ignore signal intensity outside of ROI and generates a new channel containing only voxels in the ROI using the D54D2 channel as the source channel. Imaris Surfaces function were utilized again to detect and 3D render D54D2 signals in this new channel, allowing amyloid plaque characterization restricted within microglia cells in 3D space.

### Mouse behavior procedures

All behavioral assays were performed in the light cycle (7am to 7pm). Mice were tested sequentially in the open field test, light-dark transition test, Y maze followed by novel object recognition test. All animals were habituated to the behavior room for 30 minutes before any behavior procedures.

#### Open Field Test

The open field test was performed in a large cubic box (40cm L* 40cm W * 35cm H) with an uncovered opening at the top. During the test, animals were placed at the center of the arena and allowed to freely explore the entire field for 10 minutes. Over this course, parameters including the movement of the tested animal, the total travel distance and the time spent at various regions in the arena were recorded.

#### Light-Dark Transition Test

The Light-Dark Transition test was performed in an equally illuminated rectangular box divided equally into two chambers by a partition with a door. The dark chamber (42cm L* 20cm W * 35cm H) and the partition door were made of opaque black Perspex walls and ceiling, while the light chamber (42cm L* 20cm W * 35cm H) was made of transparent Perspex walls and ceiling. A tested mouse was first placed in the dark chamber for 10 seconds with the partition door closed, and then allowed to freely explore the entire arena (light chamber and dark chamber), with the open partition door for 10 minutes. The entire travel distance and time spent in each chamber were recorded.

#### Spontaneous alternation Y maze test

The Y maze used in this assay contains three identical arms at 120 degrees from each other. Each animal was introduced to the center of the maze and was allowed to explore freely in the maze for 8 minutes. The overall number of arm entries and alterations were recorded. In this assay, the alteration percentage was defined as: Number of triads containing entries into all three arms/(total number of arm entries-2)*100.

#### Novel Object Recognition Test

The Novel Object Recognition Test was performed in a white opaque Perspex box with an uncovered top (40cm L* 40cm W * 35cm H). This test was divided into three phases (4 days): the habituation phase (day 1), the training phase (day 2 and day 3) and the testing phase (day 4). Experiments on each day started at around the same time over the day. During habituation (day 1), the tested mouse was placed at the center of box and allowed to freely explore the entire box for 10 minutes. On the following learning day (day 2), two symmetrical objects (familiar objects) with identical shape, texture, brightness and color were introduced to the box and positioned at two adjacent corners. A mouse was then placed at the center of the box and allowed to freely explore the entire box for 10 minutes. Same procedures from day 2 were repeated on day 3. On the testing day (day 4), one of the familiar objects was replaced by a novel object with different shape, texture, brightness and color. A tested mouse was placed at the center of the box and allowed to freely explore the entire box for 10 minutes. Parameters including the overall exploration time in the box, time spent on the familiar object (*T*_*familiar*_) and time spent on the novel object (*T*_*novel*_) were all recorded on the testing day. Discrimination index was calculated as DI= *T*_*novel*_ /(*T*_*novel*_ + *T*_*familiar*_).

